# A bird’s eye view to the homeostatic, Alzheimer and Glioblastoma attractors

**DOI:** 10.1101/2023.11.23.568350

**Authors:** Joan Nieves, Gabriel Gil, Augusto Gonzalez

## Abstract

Available data for white matter of the brain allows to locate the normal (homeostatic), Glioblastoma and Alzheimer’s disease attractors in gene expression space and to identify paths related to transitions like carcinogenesis or Alzheimer’s disease onset. A predefined path for aging is also apparent, which is consistent with the hypothesis of programmatic aging. In addition, reasonable assumptions about the relative strengths of attractors allow to draw a schematic landscape of fitness: a Wright’s diagram. These simple diagrams reproduce known relations between aging, Glioblastoma and Alzheimer’s disease, and rise interesting questions like the possible connection between programmatic aging and Glioblastoma in this tissue. We anticipate that similar multiple diagrams in other tissues could be useful in the understanding of the biology of apparently unrelated diseases or disorders, and in the discovery of unexpected clues for their treatment.

**Graphical abstract:** 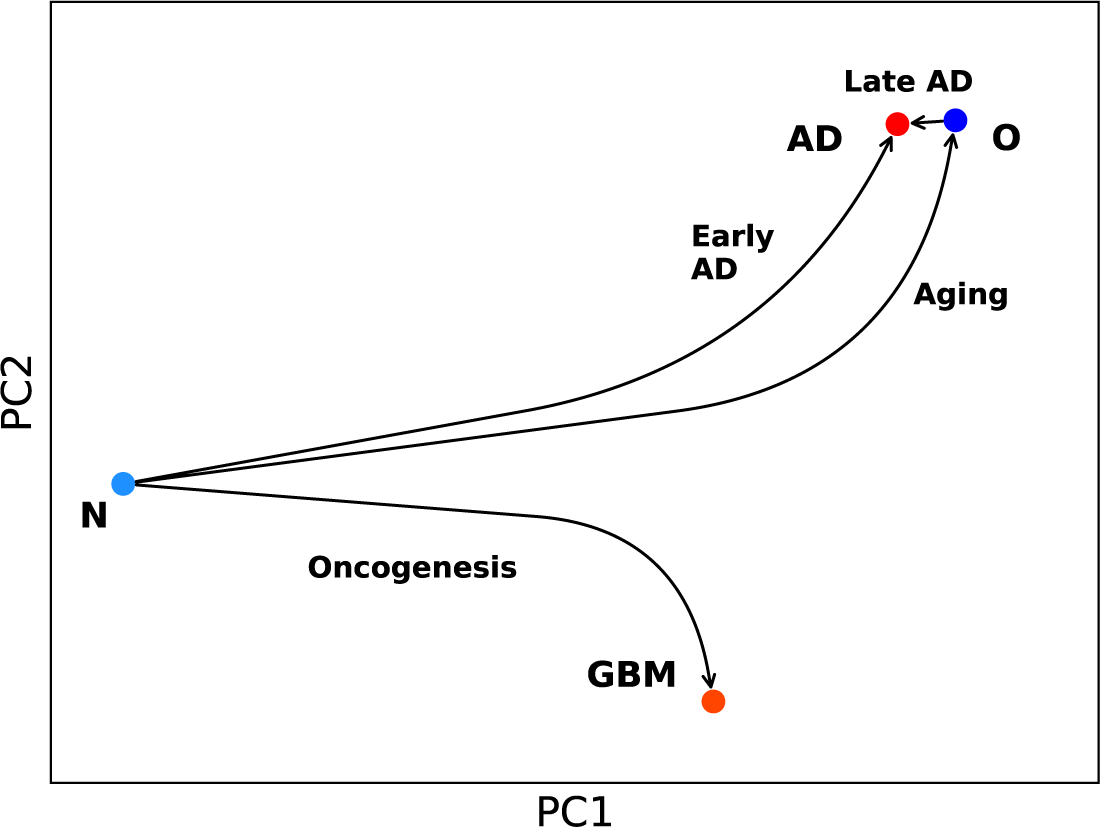

**In brief:** Aging, carcinogenesis and Alzheimer’s disease onset in white matter of the brain are shown as paths or directions in gene-expression space, a simple view that allows the analysis of their mutual relations and to rise interesting questions such as whether programmatic aging could be related to avoiding the Glioblastoma.

**Highlights:**

- Normal homeostatic, Glioblastoma and Alzheimer’s disease attractors are apparent in gene-expression space
- The relative disposition of paths for carcinogenesis and Alzheimer’s disease onset reproduce known relations between these diseases
- The observed corridor for aging is consistent with programmatic aging
- Avoiding the fall into the huge basin of the Glioblastoma could be the subject of selection pressure
- Aged normal samples could be captured by the weak Alzheimer’s disease attractor

## INTRODUCTION

A well known paradigm in molecular genetics expresses that local maxima of fitness in gene expression space are related to biological viable states [1]. This picture has been applied to the description of cell fates along differentiation lines [2]. However, to the best of our knowledge, there are no plots based on real data for a given tissue representing at least a partial landscape with more than two of these maxima. In the present paper, we provide a drawing for white matter of the brain in which the normal state (N) is represented along with the glioblastoma (GBM) attractor and the seemingly modest maximum related to Alzheimer’s disease (AD).

The plot shows that aging is a common risk factor for GBM and AD and, at the same time, that GBM and AD are opposite alternatives, as epidemiological [3,4,5,6] and molecular biology studies [7,8,9] suggest. The plot indicates also a path or corridor for normal aging, in accordance with the programmatic aging theory [10,11].

At the gene level, there are genes varying in the same way in the aging, AD progression and cancer processes, whereas there are also genes indicating the disjunctive between AD and GBM. An example of the latter is the MMP9 protein-coding gene, playing an important role in tumor invasion [12,13], but known also as a neuroprotector, controlling the interactions between axons and beta-amyloid fibers [14]. Deviations of the gene expression value from its reference in normal tissue may indicate either a potential progression to AD (under-expression) or to GBM (over-expression).

This unusual view, following from a simple plot, may help understand the relations between AD and GBM biology and identify useful gene markers for both processes. As an extra bonus, the plot allows to rise very interesting questions which are to be discussed below.

### The N + GBM + AD diagram

Our starting point is the principal component analysis [15] diagram of gene expression data for white matter of the brain, shown in **Fig. 1a**). Four groups of samples are apparent in this figure. Samples labeled as N and GBM correspond, respectively, to pathologically normal and tumor specimens in The Cancer Genome Atlas data for Glioblastoma (TCGA, https://www.cancer.gov/tcga) [16]. They are taken during surgery procedures. Tumors are geographically localized in different brain zones but, as it is common for Glioblastoma, they are white matter tumors [17]. The centers of the N and GBM clouds of samples in gene expression space define, respectively, the Normal (homeostatic) and Glioblastoma Kaufmann attractors [18,19].

**Fig. 1.**
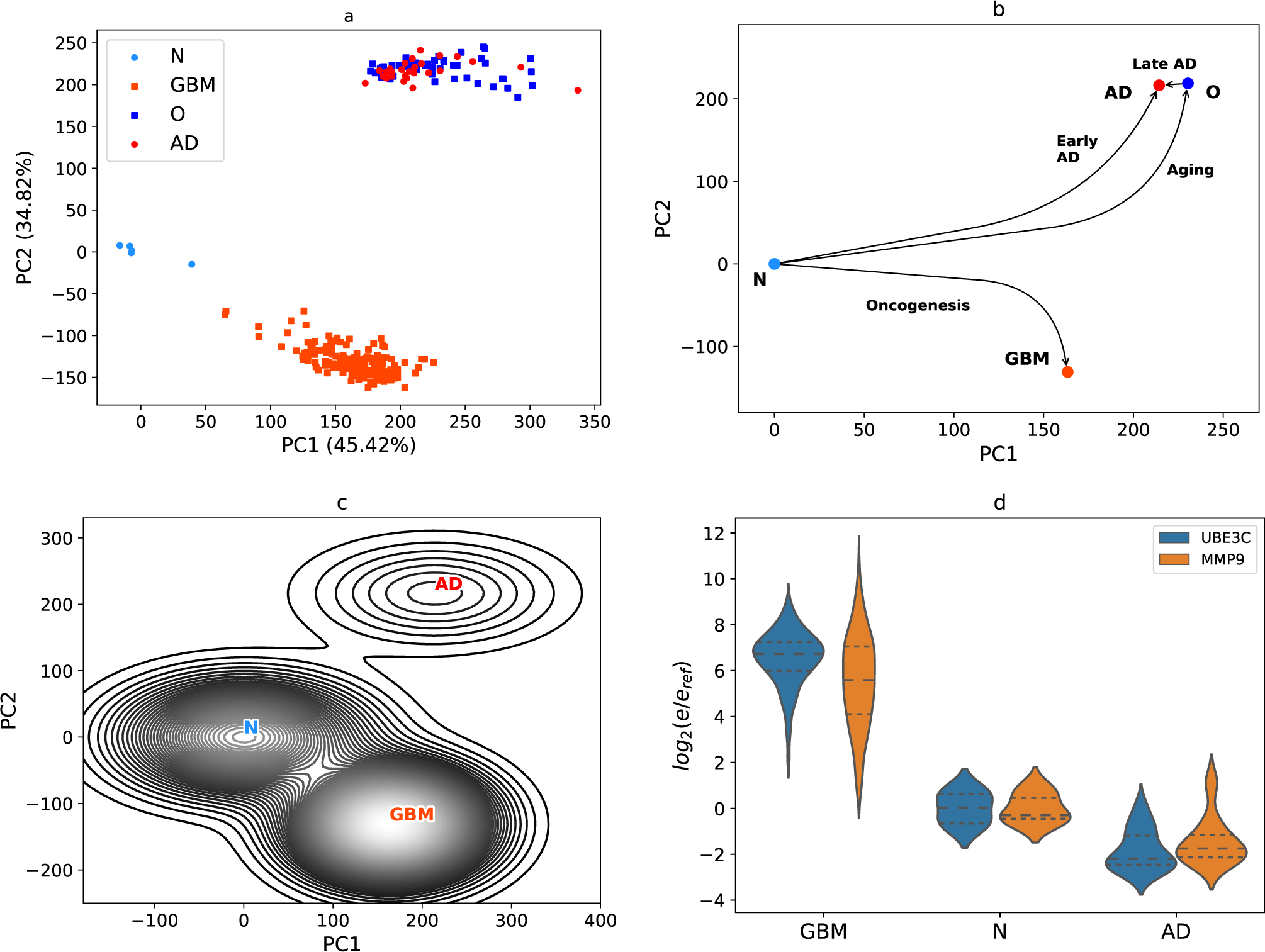
Gene expression diagrams and schematic fitness landscape. **a)** Principal component analysis of the data studied in the paper. N – Normal homeostatic state, GBM – Glioblastoma, AD – Alzheimer disease state. The normal old samples are denoted by O. **b)** Schematics of the transitions between attractors. **c)** A Wright diagram showing a hypothetical contour plot of fitness. The absolute maximum corresponds to the GBM state. The AD attractor is represented as a slight local maximum. **d)** Violin plot for the log-fold changes of MMP9 and UBE2C genes in N, AD and GBM states. The geometric mean of the expression in the N state is taken as reference in order to compute differential expression values.

On the other hand, the groups labeled as AD and O correspond, respectively, to Alzheimer disease and control white matter samples in the Allen Institute study on aging and dementia (http://aging.brain-map.org/) [20]. They are taken post mortem. The O group comes from normal aged patients, with ages ranging in the interval between 77 and 101 years. We studied the O to AD transition in Ref. [21]. As age increases, we observe a displacement of O samples towards the center of the AD cloud. Average positions of AD subgroups of samples, however, are fixed irrespective of age. This property is apparent in Fig. 4 of Ref. [21]. From these facts, we conclude that the center of the AD cloud of samples may define an attractor in gene expression space, whereas O samples are captured by the AD attractor in the process of aging.

As mentioned, we use gene expression data, in FPKM format, from Refs. [16,20]. The data was obtained by using different platforms. We took the approximately 30,000 genes that are perfectly identified in both platforms and perform a simple Principal Component Analysis (PCA) [15], described elsewhere [19,21]. The common reference used to define log-fold differential expression values and compute the covariance matrix for the PCA is the geometric mean in the N state. The 5 samples in the N state come from the TCGA data. There are also 169 GBM samples. On the other hand, in the Allen Institute data for white matter of the brain there are 47 control samples, which conform our old (O) group, and 28 AD samples.

There are both conceptual and technical issues arising when using these two dissimilar experiments in a single PCA calculation. For example, the reference N is not precisely the normal state, but a set of pathologically normal samples taken from individuals with GBM tumors. Two of the patients are even older than 70 years. From the computational side, on the other hand, one could use batch corrections [22,23], which partially amend the biases associated to each group of samples, but may introduce also uncontrolled artifacts.

Thus, we decided to take the data as it is, and use the simplest PCA technique, without any sophistication. We don’t believe that any correction will essentially change the qualitative analysis following from the 3-attractors diagrams shown in **Fig. 1a**).

The ideal situation would be to repeat the studies within a unique technological framework, and to include data from young normal people, which should be used to set the reference for differential gene expression calculations, to include data from GBM and AD patients, and data for normal patients in different age ranges. This is particularly feasible in a mouse model [24]. We look at our **Fig 1a**) diagram as a qualitative approximation to this ideal experiment.

Thus, in our approximation we get a gene expression space landscape with 3 attractors: N, GBM and AD, and a set of O samples moving towards the latter. The relative positions and main transitions between attractors are summarized in **Fig. 1b**). We assume that they are determined by the Biology underlying the processes in the tissue. The N to AD transition is labeled as “early onset of AD” in order to stress that there is also a way to AD through aging, the “late onset of AD”. A path for aging is also signaled in the figure. We shall come back to this point below.

### Fitness landscape

There is still an additional qualitative information which can be introduced in our description. It is related to a fitness variable, in such a way that we draw a kind of Wright’s diagram [1]. A schematic drawing containing a contour plot of fitness is represented in **Fig. 1c**). The N and GBM attractors are fitness maxima, and they should be separated by a low-fitness barrier [21]. The GBM should be the highest maximum [21,25]. On the other hand, the transition from O to AD is quasi-continuous, with a relatively small number of differentially expressed genes [21]. It means that there is a very small barrier or even a barrier-free path connecting O and AD. We expect a low-fitness barrier preventing the direct transitions from N to AD, and a small AD maximum, as this attractor is located in the far from N low-fitness region. All of these facts are represented in **Fig. 1c**). The scheme is constructed from a sum of Gaussians centered at the attractors, with standard deviations proportional to the actual values observed in **Fig. 1a**), and with heights qualitatively respecting the relative strengths of attractors.

Let us stress the meaning of a Wright’s diagram in a brain tissue. In other tissues somatic evolution is mainly related to stem cell replications. But, in its normal state, brain is a very slowly replicating tissue [26]. Changes in small brain regions, that is displacements in the diagram, are basically accumulated damages, i.e. aging [27]. However, once the transition to the GBM state occurs, there is an enormous increase of the replication rate of tumor cells. Let us, additionally, notice that changes related to aging are strongly apparent in white matter [28].

## MAIN RESULTS

On the basis of our diagrams, we may formulate the following remarks or statements, which are the main results of the paper:

### 1. There is a direction in gene expression space, which roughly speaking may be identified with the PC1 axis, associated with aging and with an increase in the risk for AD and GBM

Indeed, displacement along this direction implies partially climbing the low-fitness barriers separating N from the AD and GBM states, and thus augmenting the risk for both AD and GBM.

It is worth looking at the main genes involved in this process. To this end, we look at the unitary vector along the PC1 axis. Genes are ranked according to their contribution to the vector. The procedure is similar to the Page Rank algorithm [29]. We used it in our previous work [19]. A list with the first 100 genes in the ranking is given in **Supplementary Table I**. Positive amplitudes defines genes which expression increases in the displacement along the positive direction of PC1, whereas negative amplitudes refer to silenced genes. These genes should simultaneously play a crucial role in aging, GBM and AD.

Of course, due to the qualitative-only value of our analysis, the genes and specially the ranking should be taken with care. Nevertheless, notice that 20 of the silenced genes are related to the Transmission across chemical synapses pathway. In **Supplementary Table II** we list the main Reactome pathways associated to these genes [30]. There are 56 annotated genes in this set. Decreased synaptic function is a known feature of the aged brain, according to the review [31]. The second main characteristic, according to this reference, is an increased immune function, which is not particularly apparent in our set of genes. Instead, we observe genes related to Neurotoxicity of clostridium toxins [32], to a decrease of mitochondria activity [33], micro RNAs shared between AD and GBM [34], etc.

### 2. There is a direction in gene expression space, which may be roughly identified with the PC2 axis, showing that AD and GBM are excluding alternatives

Indeed, AD and GBM seem to be in opposite semiplanes. Clinical evidence [3,4,5,6] and molecular biology studies [7,8,9] support this disjunctive. Consequently, the PC2 axis involve genes inversely deregulated in AD and GBM. In **Supplementary Table III**, we list the top 100 genes defined by the unitary vector along the PC2 axis. Positive weights correspond to genes which expression increases in the N to AD transition. On the other hand, negative amplitudes correspond to genes with increasing expression in the N to GBM transition.

Only 18 of the genes in our set are annotated in Reactome pathways. The pathways are seen in **Supplementary Table IV**. They are related to control of the cell cycle, DNA replication, apoptosis, modification of the extracellular matrix, etc, i.e. to cancer hallmarks [35–37].

Above, we mentioned MMP9 as an example of genes playing opposite roles in GBM and AD. The UBE2C protein-coding gene is another known gene with this characteristic [38,39]. **Fig. 1d**) shows violin plots for the differential expression of both genes in N, AD and GBM samples. They are over-expressed in the N to GBM transition, but silenced in the early N to AD transition.

Notice also in **Supplementary Table III** the presence of many ribosome proteins, small nuclear, micro RNA and other genes, inversely regulated in both processes.

### 3. There is an aging corridor, that is a preferential path for aging in gene expression space

In our data, there are samples in the N region and samples corresponding to normal aged brains, located in a definite region close to the AD attractor. In other words, the process of aging seems to define a trajectory or corridor of continuously decreasing fitness, from which the O data shows the last segment. Samples in the intermediate region are, however, lacking.

Instead of including additional samples to our figure, which would introduce additional batch effects, we use recent results in a mouse model [24] showing undoubtedly a continuous corridor for aging. We give in **Supplementary Fig. 1 a** replot of their data for corpus callosum, a white matter rich region. In the left panel, the first two principal components are plotted for the centers of the subgroups of samples. Mouse ages between 3 and 28 months are considered, the latter is roughly equivalent to 80 years in a human scale. A corridor for aging is apparent. The right panel, on the other hand, shows true distances including all the components. Thus, the projections into the (PC1, PC2) plane are a fair representation of the actual distribution of points.

In our scheme, **Fig. 1b**), an aging corridor is delineated. **Fig. 1c**) suggests that the corridor is a direction with minimal decrease of fitness.

A preferential direction or corridor for aging is consistent with the hypothesis of programmatic aging [10,11], i.e. the idea that aging is programmed in our genes.

### 4. The predetermined aging corridor could be related to the pressure of avoiding the strong GBM attractor

A very interesting question to answer is why is it a preferred direction for aging selected. Our oversimplified scheme **Fig. 1c**) offers an unexpected answer to this question: in white matter it could be related to the pressure of avoiding the strongest GBM attractor.

Indeed, for each small portion of the tissue, we may model aging as a kind of random motion starting in the N region. A similar model was used in Ref. [40] in order to describe somatic evolution to cancer. We first assume that the direction of jumps is random in the plane shown in **Fig. 1c**). Then, there is a relatively high probability for trajectories to be captured by the huge basin of the GBM attractor leading to the initiation of a tumor. This implies an enormous increase of fitness, the spread of the tumor in brain and a life expectancy for the individual of only around two years after initiation [41]. It may impact on individuals of the reproductive age. Thus, avoiding the GBM attractor could be the subject of selection pressure.

As an indirect check, we may compare GBM and AD incidences. In a model where the direction of jumps is random, the incidence of GBM should be much higher than that of AD. However, global incidence for glioblastoma is less than 10 in 100,000 people [42], as contrasted with the 5% of AD for people in the age interval 65-74 years, and 13% of people age 75 to 84 [43]. Motion towards the GBM center is avoided.

### 5. The late onset of AD could the result of capture by the AD attractor of aged brain micro states

The picture is, thus, as follows. The process of aging is initially related to a displacement along the aging corridor with the corresponding decrease of fitness. In the last steps, the O states are captured by the weak AD attractor. This statement is supported by calculations in Ref. [21]. We already mentioned that, as a function of age, subgroups of O samples move towards the AD center.

In **Supplementary Table V** we show the top 10 genes in the O to AD transition. They involve genes included in the **Supplementary Table I**, but varying in the opposite direction, that is in the negative direction of the PC1 axis. This fact is represented in the schematic diagram given in **Fig. 1b**).

## DISCUSSION

Our simple qualitative drawings identify directions in gene expression space associated to different biological processes: aging, carcinogenesis, AD onset. Everyone of these directions is characterized by a “metagene” or gene expression profile, from which the main genes contributing to the process can be extracted.

Some of our results confirm previous knowledge, but others require further corroborations. For example, the idea that programmatic aging could be related to avoiding the strongest GBM attractor, or the late onset of AD as the capture by the AD attractor of normal aged samples. We hope, they will motivate experimental research work along these directions. Particularly feasible is a mouse model, of which Ref. [24] is a nice example.

Let us stress that even more refined data or computational methods could not essentially modify our qualitative schemes with only 3 attractors. Their relative positions could vary, but the formulated statements will remain.

**Fig. 1a**) should be completed with data corresponding to other kinds of dementia or brain disorders. In particular, one should expect a Parkinson disease area close to the AD attractor and opposite to GBM [44]. The whole picture may reveal a still finer topology of gene expression space and a richer Wright diagram.

We anticipate that similar diagrams in other tissues, besides providing an integral perspective, could be useful in the understanding of the biology of apparently unrelated diseases or disorders, and in the discovery of unexpected clues for their treatment.

### Limitations of the study

The main limitation was already mentioned: data is insufficient and imperfect. We, nevertheless, strongly believe that conclusions deriving from our qualitative three-attractors diagrams are robust against data improvement. Additional testing in a mouse model, along the same lines of the experiment shown in Ref. [24], is feasible and highly desirable.

## AVAILABILITY OF DATA AND CODE

The data we used, the Python routines and the main results are integrated in the following public GitHub repository: https://github.com/JoanANievesCuadrado/GBM-ALZ

## SUPPLEMENTAL INFORMATION

Supplemental information can be found at …

## ACKNOWLEDGMENTS

Authors acknowledge the Office of External Activities of the Abdus Salam Centre for Theoretical Physics for support. The research is carried on under a project of the Nuclear Energy and Advanced Technologies Agency (AENTA), Cuba. Authors are grateful to C. Carricarte for a critical reading of the manuscript.

## AUTHOR CONTRIBUTIONS

A.G. conceived and coordinated the work. G.G. suggested the connection with programmatic aging. J.N. processed the data and is in charge of the GitHub repository. All authors analyzed and interpreted the results, contributed to the manuscript and approved the final version.

## COMPETING INTERESTS

The authors declare that they have no competing interests.

## Supplemental Information

**Supplementary Fig. 1.**
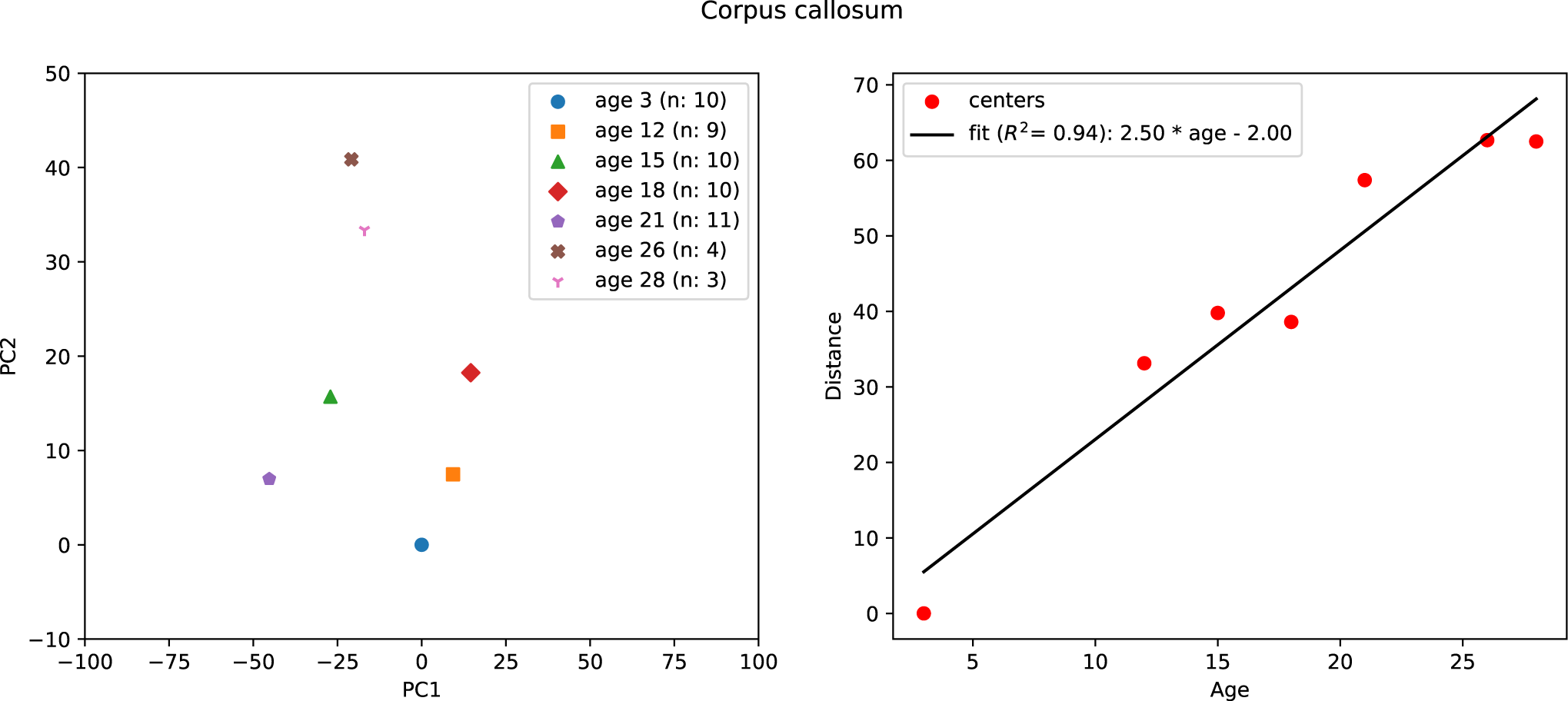
Replot of the data of Ref. [24] in a mouse model for corpus callosum, a white matter rich region. Left panel: Principal component analysis of the data. The centers of the subgroups of samples are shown. Ages between 3 and 28 months are considered. An aging direction is apparent. Right panel: Full distances (including all components) to the initial point (3 months). This figure shows that the projection to the (PC1, PC2) plane is a fair representation.

**Supplementary Table I.** The top 100 relevant genes contributing to the unitary vector along PC1 in the common plot.

| Gene | Weight in PC1 |
| --- | --- |
| RN7SL1 | 0.04 |
| RPPH1 | 0.03 |
| RNU2-1 | 0.03 |
| SNORD3A | 0.03 |
| RNU1-2 | 0.03 |
| RMRP | 0.03 |
| RNU1-4 | 0.03 |
| RNU1-3 | 0.03 |
| RNVU1-18 | 0.03 |
| RNU1-1 | 0.03 |
| SNORA73A | 0.02 |
| RNVU1-7 | 0.02 |
| RNU2-2P | 0.02 |
| WARS2-IT1 | 0.02 |
| ... | ... |
| VGF | -0.02 |
| SLC12A5 | -0.02 |
| HIPK4 | -0.02 |
| SSTR1 | -0.02 |
| CPLX1 | -0.02 |
| SYT5 | -0.02 |
| MIR657 | -0.02 |
| INA | -0.02 |
| CCK | -0.02 |
| GABRD | -0.02 |
| MIR1249 | -0.02 |
| WIF1 | -0.02 |
| NECAB2 | -0.02 |
| FBXL16 | -0.02 |
| GDA | -0.02 |
| KIAA1045 | -0.02 |
| SYNPR | -0.02 |
| HMP19 | -0.02 |
| CHGA | -0.02 |
| SNCG | -0.02 |
| SVOP | -0.02 |
| LRTM2 | -0.02 |
| WNT10B | -0.02 |
| MTND4P12 | -0.02 |
| PHYHIP | -0.02 |
| RAB3A | -0.02 |
| RASAL1 | -0.02 |
| HPCA | -0.03 |
| SERTM1 | -0.03 |
| GABRG2 | -0.03 |
| CAMKV | -0.03 |
| CREG2 | -0.03 |
| SNAP25 | -0.03 |
| GAD2 | -0.03 |
| SYT13 | -0.03 |
| KLK7 | -0.03 |
| WBSCR17 | -0.03 |
| SV2B | -0.03 |
| SLC30A3 | -0.03 |
| ATP1A3 | -0.03 |
| CPNE6 | -0.03 |
| CAMK2A | -0.03 |
| SLC6A7 | -0.03 |
| GABRA1 | -0.03 |
| KCNJ4 | -0.03 |
| CDK5R2 | -0.03 |
| CALY | -0.03 |
| GNG3 | -0.03 |
| CALB2 | -0.03 |
| CHD5 | -0.03 |
| VSNL1 | -0.03 |
| CRYM | -0.03 |
| CCKBR | -0.03 |
| NEUROD6 | -0.03 |
| HRH3 | -0.03 |
| SYT1 | -0.03 |
| SNCB | -0.03 |
| CHRM1 | -0.03 |
| MAL2 | -0.03 |
| SNORD113-3 | -0.03 |
| PNMA5 | -0.03 |
| CPLX2 | -0.03 |
| TMEM130 | -0.03 |
| LOC105373377 | -0.03 |
| SYN2 | -0.03 |
| SLC6A17 | -0.03 |
| ICAM5 | -0.03 |
| NEFL | -0.03 |
| SYN1 | -0.03 |
| C1QL3 | -0.03 |
| NEFM | -0.03 |
| DDN | -0.03 |
| FAM163B | -0.03 |
| GABRA5 | -0.03 |
| NRGN | -0.03 |
| SLC32A1 | -0.03 |
| SLC8A2 | -0.03 |
| PACSIN1 | -0.03 |
| SULT4A1 | -0.03 |
| CACNG3 | -0.03 |
| GRIN1 | -0.03 |
| SLC17A7 | -0.03 |
| PDYN | -0.03 |
| PRKCG | -0.03 |
| SST | -0.04 |
| MIR770 | -0.04 |

**Supplementary Table II.** The main Reactome pathways related to the 100 first genes in the ranking along the PC1 direction.

| Pathway name | Entities |  |  |  | Reactions |  |
| --- | --- | --- | --- | --- | --- | --- |
|  | found | ratio | p-value | FDR* | found | ratio |
| Transmission across Chemical Synapses | 20 / 344 | 0.022 | 3.72e-13 | 9.16e-11 | 63 / 167 | 0.012 |
| Neuronal System | 20 / 490 | 0.032 | 1.97e-10 | 2.42e-08 | 72 / 221 | 0.015 |

**Supplementary Table III.** The top 100 relevant genes contributing to the unitary vector along PC2 in the common plot.

| Gene | Weight in PC2 |
| --- | --- |
| RN7SL1 | 0.05 |
| RPPH1 | 0.04 |
| RNU2-1 | 0.04 |
| SNORD3A | 0.04 |
| RMRP | 0.04 |
| RNU1-3 | 0.03 |
| RNU1-1 | 0.03 |
| RNU1-4 | 0.03 |
| RNVU1-18 | 0.03 |
| RNU1-2 | 0.03 |
| SNORA73A | 0.03 |
| RNU2-2P | 0.03 |
| BCYRN1 | 0.03 |
| LOC101928075 | 0.03 |
| RNY3 | 0.03 |
| SNORA63 | 0.03 |
| RNVU1-7 | 0.03 |
| SCARNA2 | 0.03 |
| SNORA48 | 0.02 |
| CYCSP30 | 0.02 |
| SCARNA10 | 0.02 |
| RN7SL2 | 0.02 |
| LOC729348 | 0.02 |
| SCARNA5 | 0.02 |
| SCARNA7 | 0.02 |
| SNORA81 | 0.02 |
| SNORD97 | 0.02 |
| DNAJC19P8 | 0.02 |
| FCF1P1 | 0.02 |
| SNORA57 | 0.02 |
| SNORD10 | 0.02 |
| SNORA49 | 0.02 |
| SNORA54 | 0.02 |
| RAB9AP2 | 0.02 |
| RAB9AP5 | 0.02 |
| RNY1 | 0.02 |
| ... | ... |
| TPT1P9 | -0.02 |
| RPL31P4 | -0.02 |
| RPL10P9 | -0.02 |
| GAPDHP65 | -0.02 |
| RPL12P4 | -0.02 |
| RPS20P14 | -0.02 |
| FAM64A | -0.02 |
| RPS2P46 | -0.02 |
| RPS3AP5 | -0.02 |
| RPL7P1 | -0.02 |
| YBX1P1 | -0.02 |
| FTH1P7 | -0.02 |
| RPL35P5 | -0.02 |
| RPL13AP25 | -0.02 |
| RPL37AP1 | -0.02 |
| AURKB | -0.02 |
| FAUP1 | -0.02 |
| MMP9 | -0.02 |
| FTLP2 | -0.02 |
| RPL7P9 | -0.02 |
| GPX1P1 | -0.02 |
| CHI3L1 | -0.02 |
| MIR621 | -0.02 |
| PI3 | -0.02 |
| RPL6P27 | -0.02 |
| IGFBP2 | -0.02 |
| BIRC5 | -0.02 |
| TPI1P1 | -0.02 |
| FTH1P8 | -0.02 |
| RPS27AP16 | -0.02 |
| RPS27AP5 | -0.02 |
| LOC105369550 | -0.02 |
| PLA2G2A | -0.02 |
| NDUFA4P1 | -0.02 |
| FTH1P2 | -0.02 |
| TMSB4XP8 | -0.02 |
| RPL26P19 | -0.02 |
| COL1A1 | -0.02 |
| COX6A1P2 | -0.02 |
| COL3A1 | -0.02 |
| RPS27P4 | -0.02 |
| SAA1 | -0.02 |
| RPL14P1 | -0.02 |
| RPS27P3 | -0.02 |
| SNRPGP10 | -0.02 |
| MT2P1 | -0.02 |
| RPS3AP26 | -0.02 |
| PBK | -0.02 |
| TOP2A | -0.02 |
| CPXM1 | -0.02 |
| MIR3682 | -0.02 |
| RPS18P12 | -0.03 |
| RPS3AP6 | -0.03 |
| SNRPGP2 | -0.03 |
| RPS7P1 | -0.03 |
| RPS15P4 | -0.03 |
| MYBL2 | -0.03 |
| RPL41P1 | -0.03 |
| RPLP0P9 | -0.03 |
| UBE2C | -0.03 |
| RPL18AP3 | -0.03 |
| RPL39P3 | -0.03 |
| FTLP3 | -0.03 |
| RPLP0P6 | -0.03 |

**Supplementary Table IV.** The main pathways related to the 100 first genes in the ranking along the PC2 direction.

| Pathway name | Entities |  |  |  | Reactions |  |
| --- | --- | --- | --- | --- | --- | --- |
|  | found | ratio | p-value | FDR* | found | ratio |
| G0 and Early G1 | 4 / 38 | 0.002 | 1.58e-04 | 0.032 | 4 / 27 | 0.002 |
| TFAP2A acts as a transcriptional repressor during retinoic acid induced cell differentiation | 2 / 9 | 5.86e-04 | 0.002 | 0.191 | 2 / 7 | 4.83e-04 |
| Interleukin-4 and Interleukin-13 signaling | 6 / 211 | 0.014 | 0.004 | 0.254 | 2 / 47 | 0.003 |
| SUMOylation of DNA replication proteins | 3 / 50 | 0.003 | 0.005 | 0.272 | 4 / 8 | 5.52e-04 |
| Transcription of E2F targets under negative control by p107 (RBL1) and p130 (RBL2) in complex with HDAC1 | 2 / 20 | 0.001 | 0.009 | 0.336 | 2 / 8 | 5.52e-04 |
| Assembly of collagen fibrils and other multimeric structures | 3 / 67 | 0.004 | 0.012 | 0.336 | 16 / 26 | 0.002 |
| Collagen degradation | 3 / 69 | 0.004 | 0.013 | 0.336 | 16 / 34 | 0.002 |
| Transcription of E2F targets under negative control by DREAM complex | 2 / 25 | 0.002 | 0.013 | 0.336 | 2 / 12 | 8.29e-04 |
| TP53 regulates transcription of several additional cell death genes whose specific roles in p53-dependent apoptosis remain uncertain | 2 / 28 | 0.002 | 0.017 | 0.355 | 2 / 19 | 0.001 |

**Supplementary Table V.** The top 10 genes in the PCA ranking for the transition from O to AD. Results are based on calculations of Ref. [21]. Weights in the unitary vector along PC1 are normalized to the highest one.

| Gene | Extended name | Weight in PC1 |
| --- | --- | --- |
| SNAP25 | Synaptosome Associated Protein 25 | 1.00 |
| VSNL1 | Visinin Like 1 | 0.90 |
| STMN2 | Stathmin 2 | 0.90 |
| ENC1 | Ectodermal-Neural Cortex 1 | 0.89 |
| NEFL | Neurofilament Light Chain | 0.89 |
| SYT1 | Synaptotagmin 1 | 0.88 |
| RGS4 | Regulator Of G Protein Signaling 4 | 0.86 |
| CHN1 | Chimerin 1 | 0.84 |
| GABRA1 | Gamma-Aminobutyric Acid Type A Receptor Subunit Alpha1 | 0.77 |
| GABRG2 | Gamma-Aminobutyric Acid Type A Receptor Subunit Gamma2 | 0.77 |

## Notes

https://github.com/JoanANievesCuadrado/GBM-ALZ

### Competing Interest Statement

The authors have declared no competing interest.

### Summary of Updates

Added supplementary material and changed the writing style in some sections

## REFERENCES

[1] Sewall Wright. The roles of mutation, inbreeding, crossbreeding and selection in evolution. Proceedings of the 6th International Congress on Genetics 1, 356–366 (1932).

[2] Casey, M. J., Stumpf, P. S. & MacArthur, B. D. Theory of cell fate. WIREs Systems Biology and Medicine 12, e1471 (2019).

[3] Ou, S.-M. et al. Does Alzheimer’s Disease Protect against Cancers? A Nationwide Population-Based Study. Neuroepidemiology 40, 42–49 (2012).

[4] Driver, J. A. et al. Inverse association between cancer and Alzheimer’s disease: results from the Framingham Heart Study. BMJ 344, e1442 (2012).

[5] Roe, C. M. et al. Cancer linked to Alzheimer disease but not vascular dementia. Neurology 74, 106– 112 (2009).

[6] Musicco, M. et al. Inverse occurrence of cancer and Alzheimer disease: A population-based incidence study. Neurology 81, 322–328 (2013).

[7] Liu, T. et al. Transcriptional signaling pathways inversely regulated in Alzheimer’s disease and glioblastoma multiform. Scientific Reports 3, 3467 (2013).

[8] Lanni, C., Masi, M., Racchi, M. & Govoni, S. Cancer and Alzheimer’s disease inverse relationship: an age-associated diverging derailment of shared pathways. Molecular Psychiatry 26, 280–295 (2020).

[9] Cai, J. et al. Exploring the inverse association of glioblastoma multiforme and Alzheimer’s disease via bioinformatics analysis. Medical Oncology 39, 182 (2022).

[10] de Magalhães, J. P. Programmatic features of aging originating in development: aging mechanisms beyond molecular damage? The FASEB Journal 26, 4821–4826 (2012).

[11] Gems, D. The hyperfunction theory: An emerging paradigm for the biology of aging. Ageing Research Reviews 74, 101557 (2022).

[12] Choe, G. et al. Active matrix metalloproteinase 9 expression is associated with primary glioblastoma subtype. Clinical Cancer Research 8, 2894–2901 (2002).

[13] Xue, Q. et al. High expression of MMP9 in glioma affects cell proliferation and is associated with patient survival rates. Oncology Letters 13, 1325–1330 (2017).

[14] Kaminari, A., Giannakas, N., Tzinia, A. & Tsilibary, E. C. Overexpression of matrix metalloproteinase-9 (MMP-9) rescues insulin-mediated impairment in the 5XFAD model of Alzheimer’s disease. Scientific Reports 7, 683 (2017).

[15] Lever, J., Krzywinski, M. & Altman, N. Principal component analysis. Nature Methods 14, 641– 642 (2017).

[16] Brennan, C. et al. The Somatic Genomic Landscape of Glioblastoma. Cell 155, 462–477 (2013).

[17] Ellingson, B. M. et al. Probabilistic Radiographic Atlas of Glioblastoma Phenotypes. American Journal of Neuroradiology 34, 533–540 (2012).

[18] Huang, S., Ernberg, I. & Kauffman, S. Cancer attractors: A systems view of tumors from a gene network dynamics and developmental perspective. Seminars in Cell & Developmental Biology 20, 869–876 (2009).

[19] Gonzalez, A., Leon, D. A., Perera, Y. & Perez, R. On the gene expression landscape of cancer. PLOS ONE 18, e0277786 (2023).

[20] Miller, J. A. et al. Neuropathological and transcriptomic characteristics of the aged brain. ELife 6, 31126 (2017).

[21] Gonzalez, A., Nieves, J., Leon, D. A., Bringas Vega, M. L. & Sosa, P. V. Gene expression rearrangements denoting changes in the biological state. Scientific Reports 11, 8470 (2021).

[22] Haghverdi, L., Lun, A., Morgan, M. et al. Batch effects in single-cell RNA-sequencing data are corrected by matching mutual nearest neighbors. Nature Biotechnology 36, 421–427 (2018).

[23] Zhang Y., Parmigiani G., Johnson W.E. *ComBat-seq*: batch effect adjustment for RNA-seq count data. NAR Genom. Bioinform. 2, lqaa078 (2020).

[24] Oliver Hahn, Aulden G. Foltz, Micaiah Atkins, et al. Atlas of the aging mouse brain reveals white matter as vulnerable foci. Cell 186, 4117–4133 (2023). 10.1016/j.cell.2023.07.027

[25] Gonzalez, A., Quintela, F., Leon, D. A., Bringas-Vega, M. L. & Valdes-Sosa, P. A. Estimating the number of available states for normal and tumor tissues in gene expression space. Biophysical Reports 2, 100053 (2022).

[26] Kirsty L. Spalding, Ratan D. Bhardwaj, Bruce A. Buchholz, et al. Retrospective Birth Dating of Cells in Humans. Cell, Vol. 122, 133–142 (2005).

[27] Schumacher, B., Pothof, J., Vijg, J. et al. The central role of DNA damage in the ageing process. Nature 592, 695–703 (2021). 10.1038/s41586-021-03307-7

[28] C.R.G. Guttmann, F. A. Jolesz, R. Kikinis, et al. White matter changes with normal aging. Neurology 50, 972–978 (1998).

[29] Duhan, N., Sharma, A. K. & Bhatia, K. K. Page Ranking Algorithms: A Survey. 2009 IEEE International Advance Computing Conference, 1530–1537 (2009). doi:10.1109/iadcc.2009.4809246.

[30] Marc Gillespie, Bijay Jassal, Ralf Stephan, et al. The reactome pathway knowledgebase. Nucleic Acids Research 50, 2022, D687–D692 (2022).

[31] Ham, S., Lee, SJ.V. Advances in transcriptome analysis of human brain aging. Exp Mol Med 52, 1787–1797 (2020).

[32] Biazzo M, Allegra M and Deidda G. Clostridioides difficile and neurological disorders: New perspectives. Frontiers in Neurosciences 16, 946601 (2022).

[33] Nuo Sun, Richard J. Youle, Toren Finkel. The Mitochondrial Basis of Aging. Molecular Cell 61, 654–666 (2016).

[34] Thomas, L., Florio, T. & Perez-Castro, C. Extracellular Vesicles Loaded miRNAs as Potential Modulators Shared Between Glioblastoma, and Parkinson’s and Alzheimer’s Diseases. Frontiers in Cellular Neuroscience 14, 590034 (2020).

[35] Hanahan, D. & Weinberg, R. A. The Hallmarks of Cancer. Cell 100, 57–70 (2000).

[36] Hanahan, D. & Weinberg, R. A. Hallmarks of Cancer: The Next Generation. Cell 144, 646–674 (2011).

[37] Hanahan, D. Hallmarks of Cancer: New Dimensions. Cancer Discovery 12, 31–46 (2022).

[38] Ma, R. et al. High expression of UBE2C is associated with the aggressive progression and poor outcome of malignant glioma. Oncology Letters 11, 2300–2304 (2016).

[39] Jaladanki, S. K., Elmas, A., Malave, G. S. & Huang, K. Genetic dependency of Alzheimer’s disease-associated genes across cells and tissue types. Scientific Reports 11, 12107 (2021).

[40] Herrero, R., Leon, D. A. & Gonzalez, A. A one-dimensional parameter-free model for carcinogenesis in gene expression space. Scientific Reports 12, 4748 (2022).

[41] Poon, M.T.C., Sudlow, C.L.M., Figueroa, J.D. et al. Longer-term (≥ 2 years) survival in patients with glioblastoma in population-based studies pre- and post-2005: a systematic review and meta-analysis. Scientific Reports 10, 11622 (2020).

[42] Ohgaki H, Kleihues P. Epidemiology and etiology of gliomas. Acta Neuropathol. 109, 93 (2005).

[43] Alzheimer’s Association. 2023 Alzheimer’s Disease Facts and Figures. Alzheimers Dement. 19, 1598. (2023).

[44] Mencke, P. et al. Bidirectional Relation Between Parkinson’s Disease and Glioblastoma Multiforme. Frontiers in Neurology 11, 898 (2020).

